# PDGFRα^+^ Mesenchymal Stromal Cells Contribute to Epithelial Lineages during Prostate Development

**DOI:** 10.1101/2024.12.02.626405

**Authors:** Dibyo Maiti, Hsin-Jung Tien, Purna A. Joshi

**Author notes:** Correspondence should be sent to P.A.J.

## Abstract

Organ development is attributed to stem/progenitor cells which self-renew and differentiate into mature cells critical for tissue form and function. The prostate is an epithelial organ that participates in the production of seminal fluid. Bipotent and unipotent stem/progenitor cells resident in prostatic epithelia are shown to support its development. While the stroma is crucial for prostate organogenesis, precise stromal cells including immature stromal subsets involved in prostate biology are unknown. Utilizing genetic reporter and lineage tracing mouse models, we identify a PDGFRα^+^ mesenchymal population in the fibromuscular stroma that harbors progenitors which undergo a mesenchymal-to-epithelial transition, generating prostatic luminal and basal epithelial lineages during postnatal development. Further, these mesenchymal progenitors and their epithelial progeny persist in the adult, demonstrating their self-renewal potential. Our findings unveil a prostatic progenitor beyond the epithelium, laying down a framework for probing the contribution of stromal progenitors to the normal and diseased prostate.

**Graphical Abstract:** 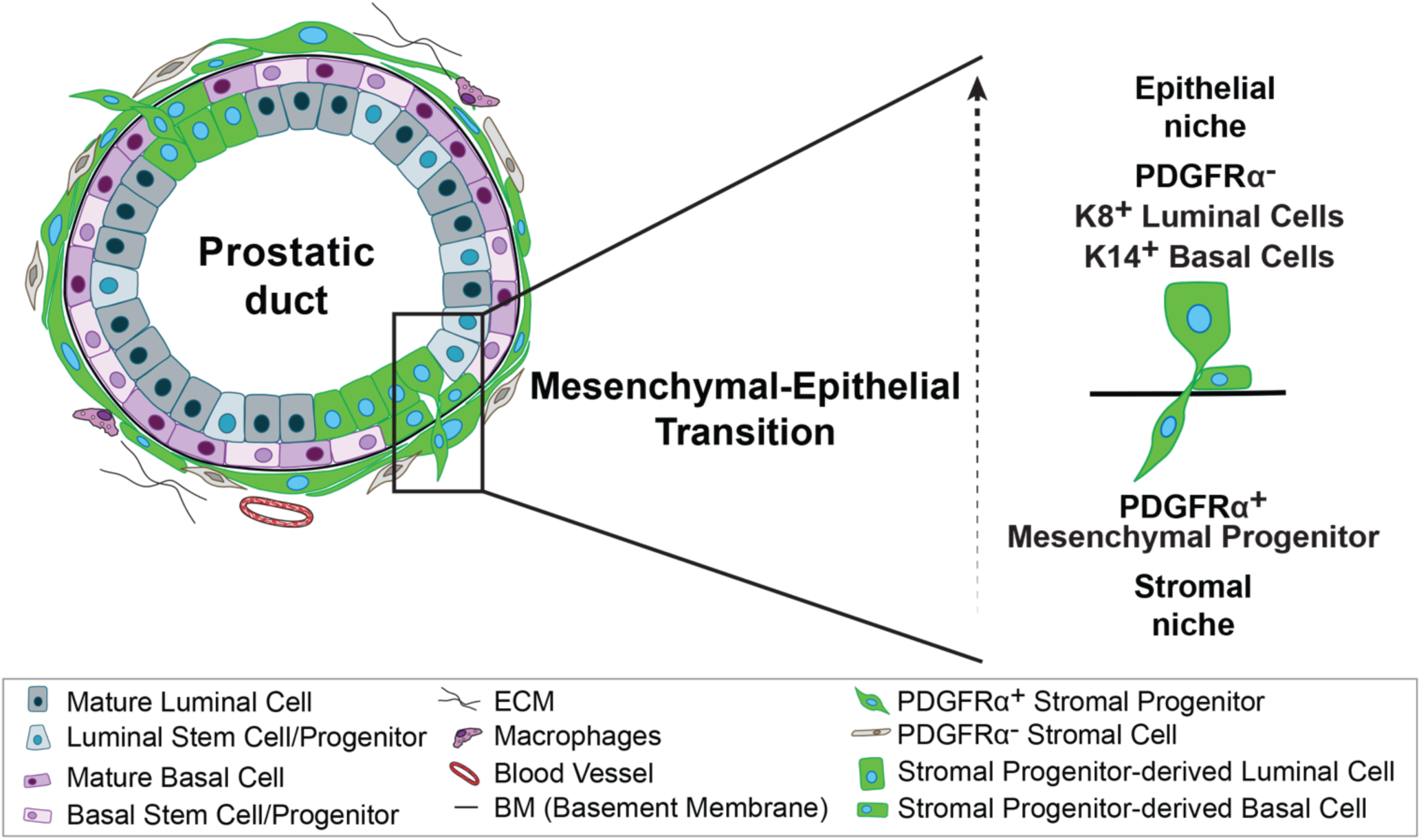

## Introduction

The prostate is a major male sex-accessory endocrine gland whose secretions contribute to seminal fluid. The organogenesis of the prostate in mice begins at embryonic day 16 and the first epithelial buds emerge at embryonic day 17.5 from the urogenital sinus (UGS)(Timms et al., 1994). In humans, prostate budding begins at 10-12-week gestation(Aaron et al., 2016). Through a series of inductive signals and lineage specification steps that are not well understood, the mouse and human prostate epithelium develops to comprise luminal, basal and rare neuroendocrine cells in adulthood(Baures et al., 2022). Development of the prostate is directed by androgens and cell-cell interactions between the mesenchyme and epithelia(Thomson and Marker, 2006). During embryonic prostate development, the androgen receptor (AR) is only expressed by the mesenchyme and not by the prostatic epithelia, and mesenchymal AR is indispensable for epithelial budding(Cunha and Chung, 1981; Lasnitzki and Mizuno, 1980). The postnatal prostate microenvironment mainly constitutes the fibromuscular prostatic stroma. This fibromuscular stroma contains mesenchymal cells such as smooth muscle cells and fibroblasts, and also blood vessels and nerves(Ittmann, 2018). Although AR is expressed by epithelial cells in the postnatal prostate, the mesenchymal stroma continues to play a major role in postnatal development and homeostasis(Cunha et al., 1983; Wei et al., 2022). The importance of the mesenchyme is further exemplified by tissue recombinant experiments performed under the kidney capsule where urogenital mesenchyme is shown to be required and inductive for prostatic epithelial growth and differentiation(Cunha and Chung, 1981; Xin et al., 2003).

As in other epithelial tissues, prostate morphogenesis is driven by tissue-resident stem and progenitor cells. Lineage tracing experiments have shown that the basal epithelial cell population contains both multipotent and unipotent progenitor cells while the luminal population has unipotent progenitors which together contribute to prostatic epithelial development(Ousset et al., 2012; Pignon et al., 2013; Tika et al., 2019). Although epithelial progenitors have been intensely studied in prostate biology, stromal progenitors emanating from the urogenital mesenchyme which generate mesenchymal cell populations present in the prostate remain elusive. Furthermore, whether such stromal progenitors could directly contribute to the prostate epithelial hierarchy has not been explored.

Here, we show that Platelet-Derived Growth Factor Receptor alpha (PDGFRα) marks a mesenchymal population that enriches for stromal progenitors in the postnatal prostate. These stromal progenitors residing adjacent to the prostatic epithelia transition to luminal and basal cells during prostate branching morphogenesis and epithelial expansion. Our study reveals PDGFRα^+^ mesenchymal stromal progenitors in the prostatic stroma as a source of epithelial descendants in the developing prostate.

## Results

### PDGFRα marks ubiquitous mesenchymal stromal cells in the developing prostate

PDGFRα has been shown to mark mesenchymal stromal cells in the endometrium, mammary gland and intestine(Brugger et al., 2020; Joshi et al., 2019; Kirkwood et al., 2022). A previous single-cell RNA sequencing study used *Pdgfrα* to cluster and identify an adult prostatic mesenchymal fibroblast population(Graham et al., 2023). However, the spatial distribution and dynamics of PDGFRα^+^ stromal cells in prostate development have not been described. To this end, we employed a *Pdgfrα^H2B-eGFP^* reporter mouse model that expresses the H2B-GFP histone fusion protein under the control of the *Pdgfrα* promoter (**Figure 1A**). GFP^+^ cells in this model will thus reflect endogenous PDGFRα expression *in vivo*. At postnatal day five (P5), we found nuclear GFP^+^ cells reporting for *Pdgfrα* expression in the prostate that colocalized with cells positive for cell surface PDGFRα by immunofluorescence and confocal imaging (**Figure 1B**). Of note, GFP^+^ cells did not express the pan-epithelial marker EpCAM and basal marker keratin 14 (K14), showing that *Pdgfrα*-expressing cells are stromally-restricted at this stage of development.

**Figure 1.**
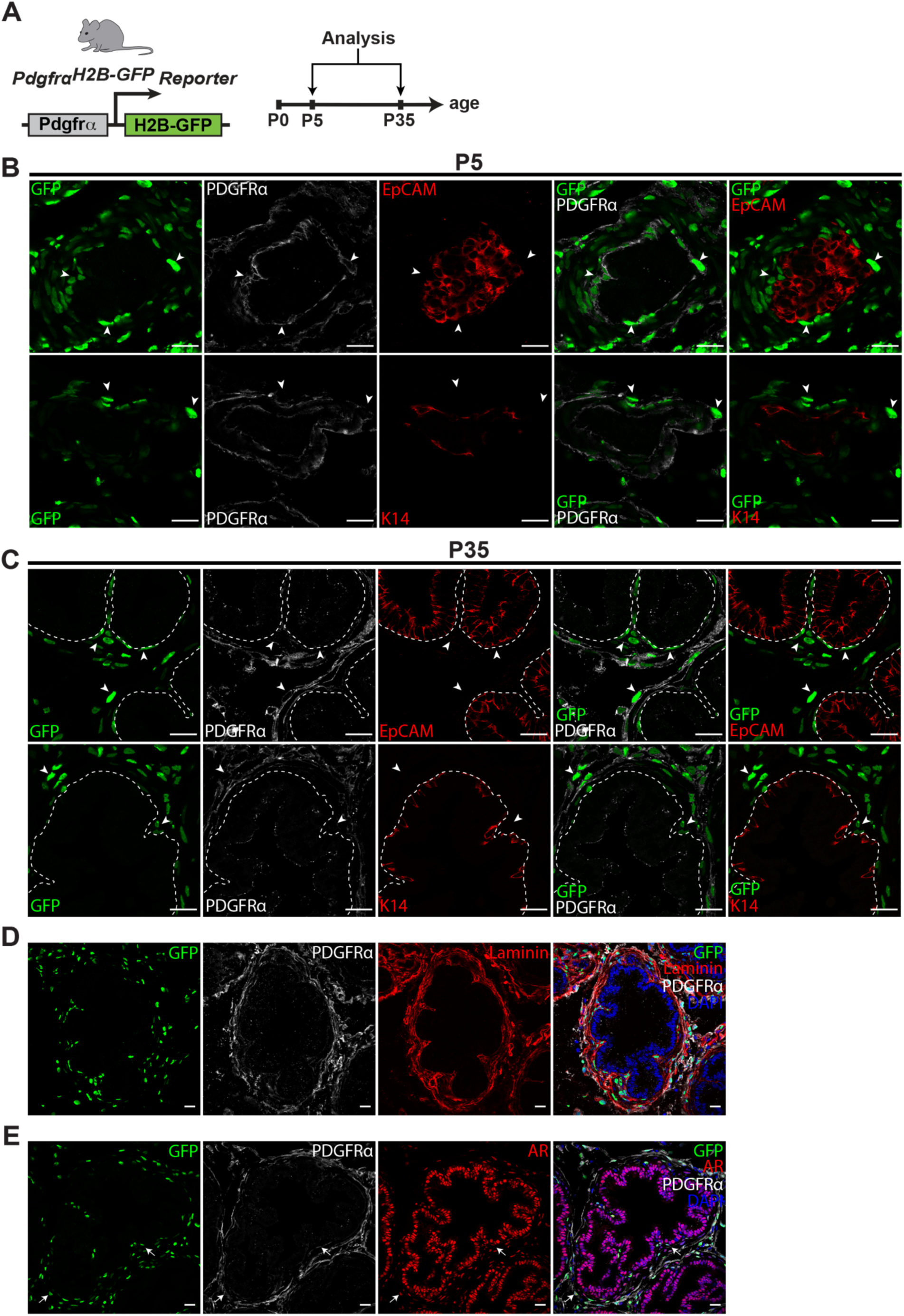
PDGFRα^+^ cells are localized to the stromal compartment in the developing postnatal mouse prostate. **(A)** Schematic illustrating the *Pdgfrα^H2B-eGFP^*reporter mouse model and postnatal timepoints of analysis. **(B)** Co-immunostaining for GFP, EpCAM, PDGFRα and GFP, K14, PDGFRα in the DLP of *Pdgfrα^H2B-eGFP^* reporter mice at P5 (representative of n=5 mice, 5 fields per tissue section); scale bars = 20 µm. Arrowheads point to GFP^+^PDGFRα^+^ cells in the prostatic stroma. **(C)** GFP, EpCAM, PDGFRα and GFP, K14, PDGFRα in the DLP of reporter mice at P35 (representative of n=6 mice, 5 fields per tissue section); scale bars = 20 µm. Arrowheads point to GFP^+^PDGFRα^+^ cells in the prostatic stroma. **(D)** GFP, Laminin, PDGFRα in the DLP of reporter mice at P35 (representative of n=3 mice, 3 fields per tissue section); scale bars = 20 µm. (**E)** GFP, AR, PDGFRα in the DLP of reporter mice at P35 (representative of n=3 mice, 3 fields per tissue section); scale bars = 20 µm. Arrows point to GFP^+^AR^+^PDGFRα^+^ cells in the prostatic stroma.

In mice, postnatal prostate development occurs from birth to 5 weeks of age with completion of branching morphogenesis by around 5 weeks of age(Sugimura et al., 1986). The murine prostate also comprises three distinct lobes which include the anterior lobe (AP), ventral lobe (VP) and dorsolateral lobe (DLP)(Timms *et al*., 1994). When mice were analyzed at 5 weeks of age (P35), we observed that GFP^+^ and PDGFRα^+^ cells remained localized to the prostatic stroma encasing EpCAM^+^ or K14^+^ epithelia in the DLP (**Figure 1C**) and also in AP and VP lobes (**Figure S1A**). These GFP^+^ mesenchymal cells were seen adjacent to the prostatic epithelium and in the interstitial space, which indicate that they are represented in spatially distinct fibroblast populations previously reported in the prostate(Peng et al., 2013) (**Figure S1B**). Co-immunostaining for the basement membrane constituent laminin confirmed the distribution of *Pdgfrα*-expressing mesenchymal cells specifically in the stroma and outside the epithelial boundary (**Figure 1D**). GFP^+^ cells also expressed nuclear AR (**Figure 1E**), suggesting likely androgen-mediated effects on these mesenchymal cells during prostate development. These data show that PDGFRα^+^ cells are vastly distributed in the mesenchymal stroma, absent in epithelia and express AR during postnatal prostate development.

### The PDGFRα^+^ mesenchymal stromal cell population exhibits progenitor activity

Since we noted widespread *Pdgfrα*-expressing cells in the stromal compartment of the developing prostate, we interrogated whether this cell population contains stromal progenitors. Based on previously established cell-surface markers that segregate prostatic cell populations(Crowell et al., 2019; Giafaglione et al., 2023), luminal (EpCAM^+^CD49f^lo^), basal (EpCAM^+^CD49f^hi^), and stromal (EpCAM^-^CD49f^-^) subsets in P35 *Pdgfrα^H2B-eGFP^*reporter mice were analyzed by flow cytometry (**Figure 2A, B**). As observed by immunostaining and confocal imaging, we were able to only detect a prominent GFP^+^ subpopulation (86 ± 2%) in the EpCAM^-^CD49f^-^ stromal subset, further supporting the localization of *Pdgfrα*-expressing cells in the prostatic stroma during development (**Figure 2B, C**). Mesenchymal progenitors in white adipose tissue depots(Merrick et al., 2019) and the mammary gland(Joshi *et al*., 2019) are known to be labeled by PDGFRα. To assess whether prostatic PDGFRα^+^ stromal cells have progenitor potential, GFP^+^ and GFP^-^ stromal cell fractions from the EpCAM^-^CD49f^-^ subset of *Pdgfrα^H2B-eGFP^* P35 prostates were isolated using Fluorescence-Activated Cell Sorting (FACS). Sorted cell fractions were plated on feeder cells in a Colony Forming Cell (CFC) assay to measure clonogenic capacity indicative of progenitor activity, as performed for other tissue progenitors(Kronstein-Wiedemann and Tonn, 2019; Stingl, 2009). We observed robust colony forming potential of GFP^+^ stromal cells whereas the GFP^-^ stromal fraction did not give rise to a single colony, showing that all the proliferative stromal progenitor activity is confined to the *Pdgfrα*-expressing population. (**Figure 2D, E**). Additionally, cell proliferation detected by Ki67 immunostaining was evident in GFP^+^ stromal cells at P5 and P35 in the developing prostate (**Figure 2F**). Thus, *Pdgfrα*-expressing cells in the prostate are proliferative and enrich for stromal progenitors.

**Figure 2.**
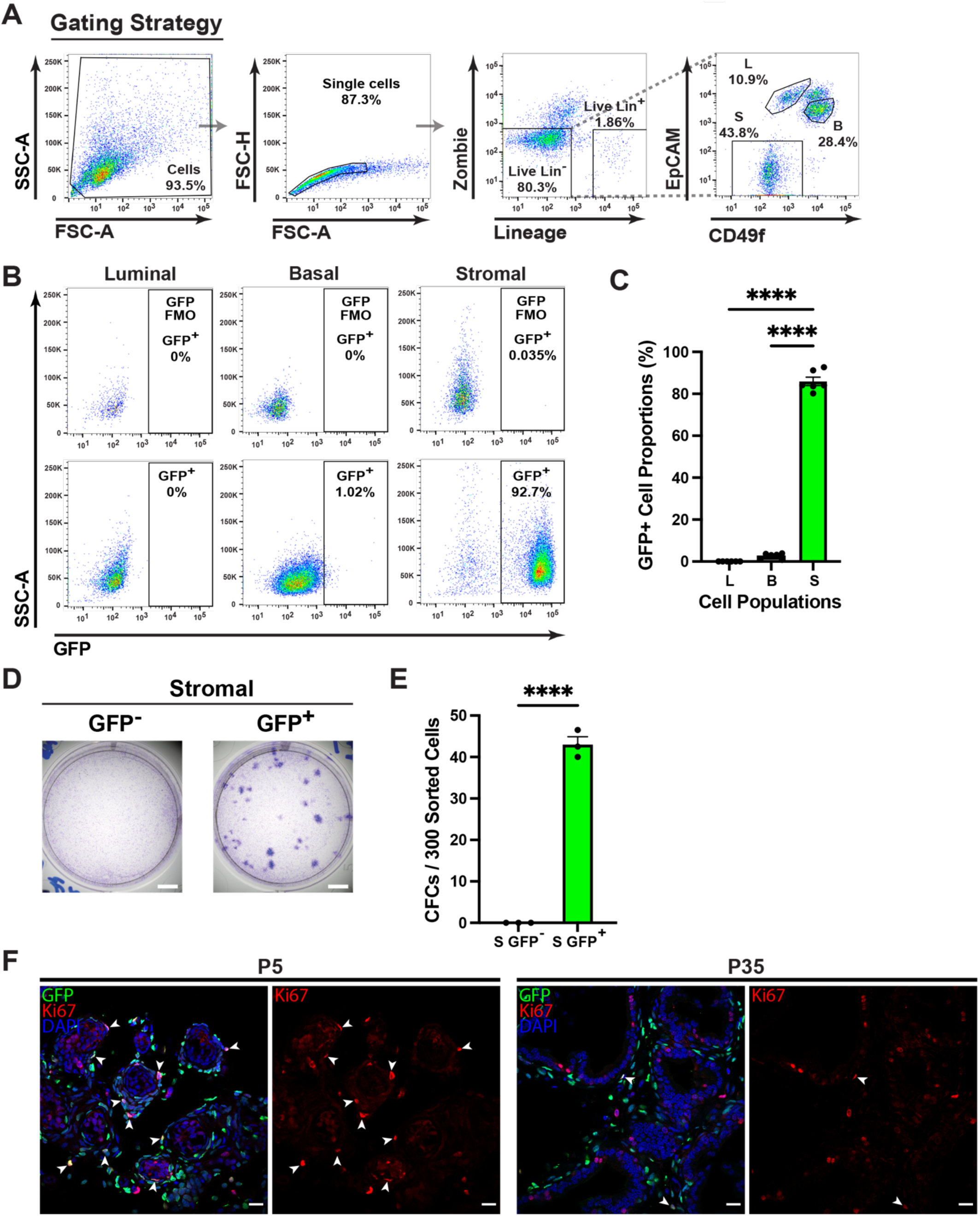
The PDGFRα^+^ mesenchymal subpopulation is enriched in stromal progenitors. **(A)** Gating strategy used in flow cytometry to exclude doublets, dead cells (Zombie UV^+^), and Lineage^+^ (Lin^+^; CD31^+^CD45^+^Ter119^+^) cells from analysis of GFP^+^ cells within the Lin^-^ population comprising prostatic stromal and epithelial subpopulations of *Pdgfrα^H2B-eGFP^*reporter P35 mice segregated using the indicated cell surface markers. **(B)** Flow cytometry analyses of GFP^+^ cells within prostatic subpopulations derived from developing prostates of P35 mice (representative of n = 5 mice); fluorescence minus one (FMO) control for GFP. **(C)** Quantification of PDGFRα^+^ cells in (B). Data represent mean ± s.e.m. ****p<0.001 (one-way ANOVA). **(D)** Representative plates of colony forming cell (CFC) assay performed on sorted GFP^+^ and GFP^-^ stromal cells (n = 3 mice); Scale bars = 5 mm. **(E)** CFC capacity of GFP^+^ and GFP^-^ stromal cells (n = 3 mice). Data represent mean ± s.e.m. ****p< 0.0001 (t-test). **(F)** Co-immunostaining for GFP, Ki67 in the DLP of *Pdgfrα^H2B-eGFP^* reporter mice at P5 and P35 (representative of n=3 mice, 5 fields per tissue section); scale bars = 20 µm. Arrowheads show GFP^+^Ki67^+^ cells in the prostatic stroma.

### PDGFRα^+^ stromal progenitors undergo MET and contribute to prostatic epithelia during early postnatal development

Next, we probed the fate of PDGFRα^+^ stromal progenitors *in situ* during postnatal prostate development using an inducible *Pdgfrα^CreERT2^R26^mTmG^* lineage tracing mouse model. In this model, PDGFRα^+^ cells are permanently marked with GFP upon tamoxifen (TAM)-induced Cre-mediated excision of a tdTomato cassette and a stop sequence, after which all subsequent progeny of the PDGFRα^+^ cell lineage will be GFP-labeled. Male mice were injected with TAM at P5 and prostates analyzed in a short trace after 24 hours to determine initially labeled cells, and then analyzed in a long trace at P35 to map GFP-labeled progeny following early postnatal prostate development (**Figure 3A**). In immunostained tissue sections of isolated prostates from P6 mice after the short trace, GFP^+^ cells were located in the prostatic stroma surrounding epithelial ducts marked by EpCAM, and these stromal GFP^+^ cells were also PDGFRα^+^ as anticipated (**Figure 3B**). Following a long trace from P5 to P35 when the bulk of branching morphogenesis is completed, GFP-labeled cells were remarkably found in prostatic epithelial ducts in addition to stroma (**Figure 3C, D**). Specifically, GFP^+^ cells in the epithelium co-expressed EpCAM but did not express PDGFRα. Prostate tissues from oil-injected *Pdgfrα^CreERT2^R26^mTmG^* control mice did not present GFP^+^ cells, precluding any leaky Cre expression (**Figure S2A, B**). When immunostained for the basal marker K14 and the luminal marker K8, GFP^+^ cells in the epithelium were observed to be K14^+^ in the basal layer and K8^+^ in the luminal layer, indicating the presence of GFP-labeled stromal progeny in both basal and luminal epithelial lineages (**Figure 3D**). Interestingly, stromal progenitor-derived GFP^+^ epithelial cells also expressed AR in addition to GFP^+^ cells in the stroma (**Figure S3**). This data demonstrate that PDGFRα^+^ stromal progenitors in the prostate undergo a mesenchymal-to-epithelial transition (MET) during early postnatal development to contribute to basal and luminal prostatic epithelial cell populations.

**Figure 3.**
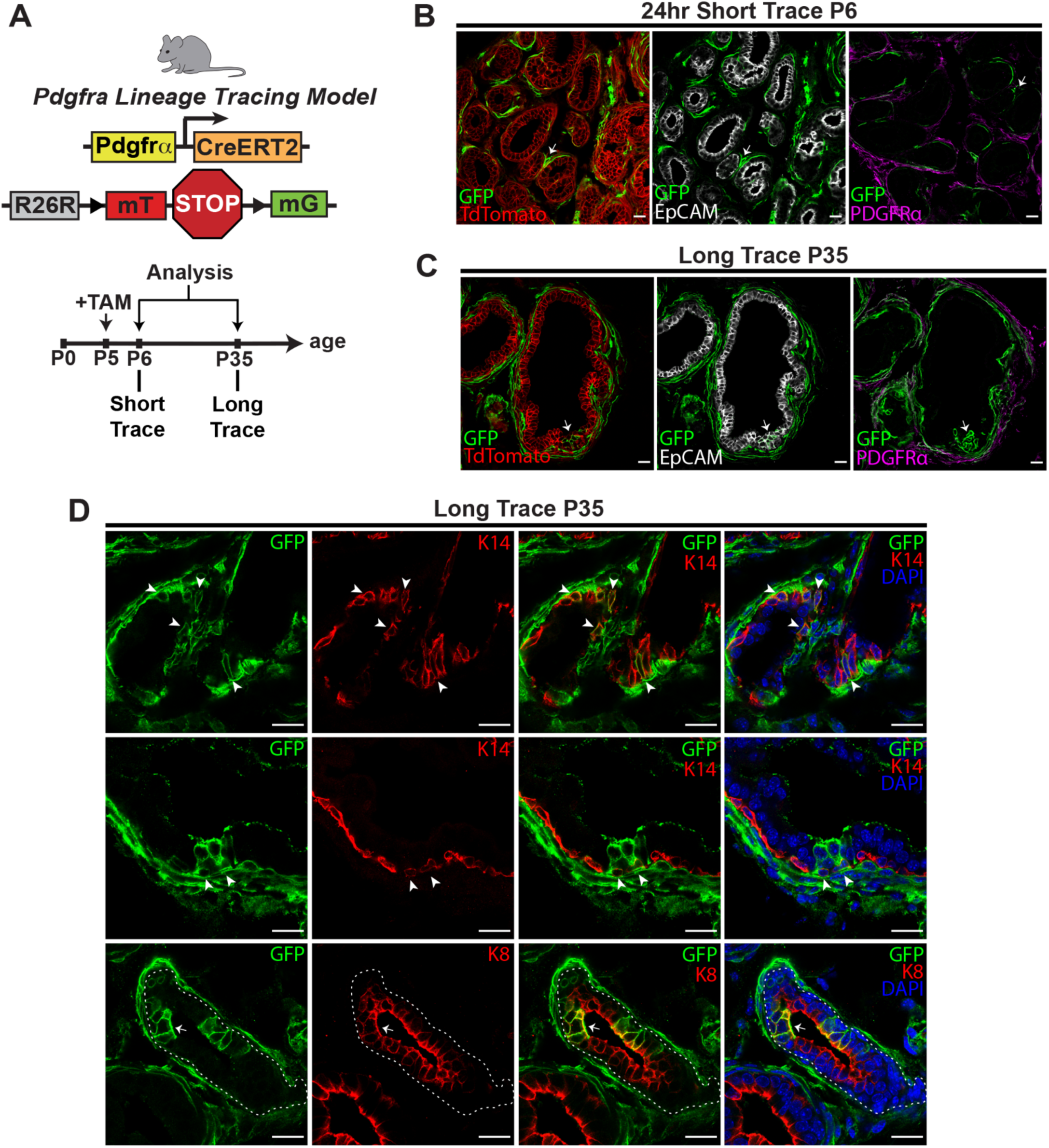
PDGFRα^+^ stromal progenitors contribute to prostatic basal and luminal cell layers during early postnatal development. **(A)** Schematic illustration of the *Pdgfrα^CreERT2^R26^mTmG^*lineage tracing mouse model and timepoints analyzed post-induction. **(B)** Co-immunostaining for GFP, tdTomato and GFP, EpCAM and GFP, PDGFRα in consecutive sections of the DLP of lineage tracing mice at P6, 24 hours after TAM induction (Short Trace) (representative of n=4 mice, 4 fields per tissue section); scale bars = 20 µm. **(C)** GFP, tdTomato and GFP, EpCAM and GFP, PDGFRα in consecutive sections of the DLP of lineage tracing mice at P35, 30 days after TAM induction (Long Trace) (representative of n=7 mice, 5 fields per tissue section); scale bars = 20 µm. Arrows in B and C point to GFP^+^ cells. **(D)** GFP, K14 and GFP, K8 in the DLP of lineage tracing mice at P35 (representative of n=5 mice, 5 fields per tissue section), scale bars = 20 µm. In D, arrowheads show GFP^+^K14^+^ cells and arrows show GFP^+^K8^+^ cells in the prostatic epithelia.

### Stromal progenitor-derived epithelial descendants persist in the adult prostate

Male mice reach sexual maturity at 6 weeks of age, but the prostate continues to slowly accumulate new branch points until puberty is complete. To determine whether PDGFRα^+^ stromal progenitor-derived epithelial cells contribute to the mature prostate, *Pdgfrα^CreERT2^R26^mTmG^*lineage tracing mice induced with TAM at P5 were analyzed at postnatal day 60 (P60) (**Figure 4A**). As seen at P35, immunostaining of prostate tissue sections at P60 showed the presence of GFP^+^PDGFRα^+^ stromal cells, and additionally, GFP^+^K14^+^ basal cells and GFP^+^K8^+^ luminal cells in the prostate epithelium which did not express PDGFRα (**Figure 4B**). Of note, GFP-labeled epithelial cells were detected in all the lobes of the prostate including the DLP, AP and VP (**Figure S4**). To quantify the dynamic contribution of stromal progenitor-derived epithelial descendants to the postnatal prostate, we enumerated EpCAM^+^GFP^+^ cells at P6 (short trace), P35 (long trace during puberty), and P60 (long trace until maturity). We found that the prostate at P60 exhibited the highest proportion of EpCAM^+^GFP^+^ cells (17.8±1.4%) compared to P35 (6.3±1%), whereas EpCAM^+^GFP^+^ cells were not detected in the short trace as stated previously (**Figure 4C, D**). We also noted Ki67^+^ cell proliferation in GFP^+^ stromal cells at P5, P35 and P60, and GFP^+^ epithelial cells at P35 and P60 (**Figure 4E).** This data infer that prostatic epithelial descendants of PDGFRα^+^ stromal progenitors self-renew, proliferate and expand into adulthood.

**Figure 4.**
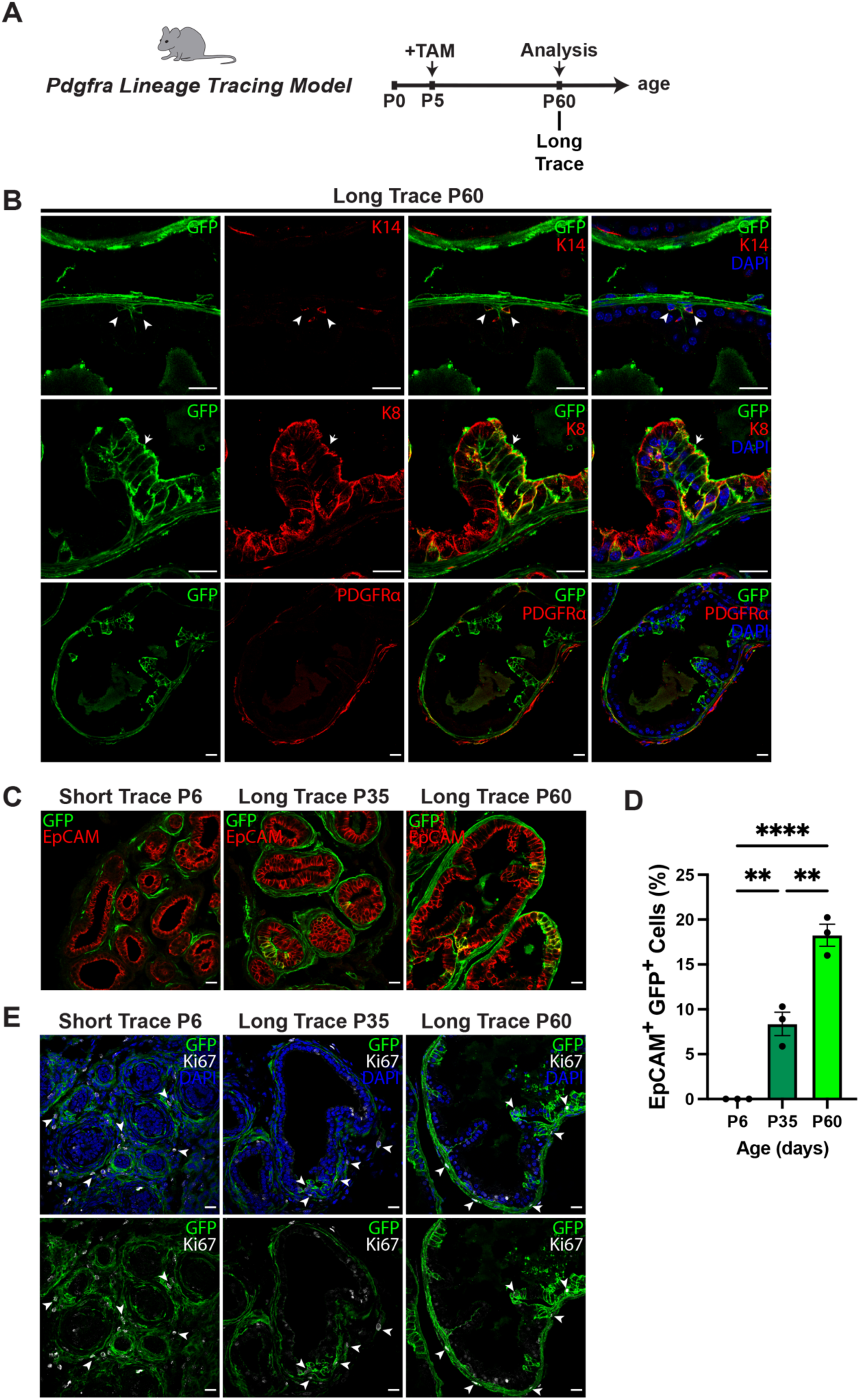
Stromal progenitors and their epithelial descendants are long-lived in the mature prostate. **(A)** Schematic of timepoint analyzed in *Pdgfrα^CreERT2^R26^mTmG^* lineage tracing mice to study stromal progenitor contribution in the adult prostate. **(B)** Co-immunostaining for GFP, K14 and GFP, K8 in the DLP of lineage tracing mice at P60; scale bars = 20 µm (representative of n=3 mice, 5 fields per tissue section). Arrowheads show GFP^+^K14^+^ cells and arrows show GFP^+^K8^+^ in mature prostatic epithelia. **(C)** GFP, EpCAM in the DLP of lineage tracing mice at P5, P35 and P60 (representative of n = 3 mice each, 3 fields per tissue section); scale bars = 20 µm. **(D)** Quantification of EpCAM^+^ GFP^+^ cells over total EpCAM^+^ in prostatic epithelia of the DLP of lineage tracing mice at P5, P35 and P60 (n = 3 mice each, 3 fields per tissue section). Data represent mean ± s.e.m. **p<0.01, ****p<0.001 (one-way ANOVA). **(E)** Co-immunostaining for GFP and Ki67 in the DLP of lineage tracing mice at P5, P35 and P60 (representative of n = 3 mice each, 3 fields per tissue section). Scale bars = 20 µm. Arrowheads show GFP^+^Ki67^+^ cells in the prostatic stroma and epithelia.

## Discussion

While the mesenchymal stromal microenvironment is known to provide critical inductive signals that govern prostatic epithelial cell fate, the existence of stromal progenitors that could directly contribute to the genesis of prostatic epithelial lineages were not previously described. Our study illuminates a PDGFRα^+^ mesenchymal progenitor that exhibits plasticity and a lineage switch to an epithelial cell fate during postnatal prostate development. Lineage tracing studies in female mice have previously documented MET in PDGFRα^+^ stromal cells which contribute to epithelial lineages in the mammary gland and uterus(Joshi *et al*., 2019; Kirkwood *et al*., 2022). In male mice, previous studies utilizing PDGFRα^+^ lineage tracing models have only shown the differentiation capacity of this lineage in fat depots to mature adipocytes(Berry and Rodeheffer, 2013; Lee et al., 2012), but not in the male reproductive tract. Our findings underscore the contribution of the PDGFRα^+^ stromal lineage to both luminal and basal prostatic epithelial cells during development, and the sustained presence of stromal progeny in the mature prostate epithelium.

Knowledge of stromal cells in prostate development has primarily been limited to the importance of AR^+^ stromal cells in promoting epithelial morphogenesis through paracrine signals(Cunha *et al*., 1983; Lasnitzki and Mizuno, 1980). Stromal cells marked by Lgr5 in proximal ducts of mouse prostatic lobes are reported to play a role in prostate tissue homeostasis(Wei *et al*., 2022). However, Lgr5^+^ stromal cells were traced in adult mice and no epithelial contribution from this stromal population was noted. Lineage tracing during postnatal prostate development has shown multipotent basal progenitors capable of differentiating into basal and luminal cells, as well as lineage-restricted basal and luminal progenitors(Ousset *et al*., 2012; Tika *et al*., 2019). Our identification of a stromal progenitor adds a new layer to the existing prostate epithelial hierarchy during development wherein mesenchymal progenitors exit their mesenchymal niche, become PDGFRa^-^ and incorporate into the epithelial niche to generate both luminal and basal epithelial descendants, thus contributing to prostate epithelia during postnatal prostate development.

Prostate development and homeostasis is highly androgen-dependent(Cunha *et al*., 1983; Xie et al., 2017). Our observation that GFP^+^PDGFRα^+^ stromal cells and their GFP^+^ epithelial progeny are AR^+^ suggest that the mesenchymal-to-epithelial cell fate transition and expansion of stromal progenitors as well as their progeny are possibly androgen-driven in the developing prostate. The transition of stromal progenitors to an epithelial cell fate in the mammary gland is also seen in response to female sex hormones estrogen and progesterone(Joshi *et al*., 2019). Intriguingly, both prostate and mammary epithelial morphogenesis also depend on inductive cues from the mesenchyme during development, and both glands are composed of luminal and basal epithelial lineages. Thus, the capacity of stromal progenitors to differentiate into epithelial populations is likely a conserved phenomenon in sex hormone-responsive organs.

Cell types and signals involved in prostate development are also implicated in prostate diseases such as cancer. Androgen signaling through AR is integral for prostate development and likewise drives prostate cancer pathogenesis(Watson et al., 2015). Previous studies have shown that both basal and luminal epithelial cells can serve as the cell-of-origin for prostate adenocarcinoma(Goldstein et al., 2010; Park et al., 2016; Stoyanova et al., 2013). Since PDGFRα^+^ stromal cells have the capacity to give rise to basal and luminal epithelial lineages in the prostate, it is interesting to speculate their potential involvement in cancer initiation or progression which warrants further investigation. The stroma has generally been ascribed a supportive role in prostate development and cancer. Our research described here uncovers, for the first time, a stromal progenitor in the prostate that has a unique potency to generate epithelial lineages, thus directly impacting prostatic epithelial cell fate. It also builds a foundation for further deciphering the dynamics and contribution of stromal progenitors to prostate biology which can be potentially tapped into for advancing prostate regenerative and cancer therapies.

## Methods

### Mice

All mice were on the C57BL/6J background and maintained on a 12 h light, 12 h dark cycle with access to food and water ad libitum. C57BL/6J wild-type (Stock# 000664), *Pdgfrα^H2B-eGFP^* (Stock# 007669), *Pdgfrα^CreERT2^* (Stock# 032770) and *Rosa26^mTmG^* (Stock# 007676) strains were procured from the Jackson Laboratory. Treatment and care of wild-type and transgenic murine models were in accordance with protocols reviewed by the Institutional Animal Care and Use Committee (IACUC) at the University of Texas at Dallas.

### Lineage Tracing

For lineage tracing studies, *Pdgfrα^CreERT2^R26^mTmG^*lineage tracing male mice were induced with a single intraperitoneal injection of 4-hydroxytamoxifen (TAM; Sigma, Cat.# H7904) in young neonatal male mice at P5 (150µg TAM diluted in 30μl sunflower seed oil) to induce Cre-mediated recombination and GFP labeling. As a control for TAM, *Pdgfrα^CreERT2^R26^mTmG^*lineage tracing male mice were injected with 30μl of sunflower seed oil and prostates were isolated for analysis at P6, P35 or P60.

### Immunostaining and confocal imaging

For immunofluorescence, freshly isolated prostates from P5 mice or microdissected individual prostate lobes from P35 and older mice were fixed in 4% paraformaldehyde at room temperature, washed with PBS, incubated in 30% sucrose overnight at 4°C and embedded in OCT after which samples were stored at -80°C and cryosectioned at 10μm thickness for immunofluorescence. Cryosections were incubated in blocking buffer (5% normal donkey serum, 1% BSA, 0.2% Triton in PBS) for 1 h at room temperature and then in primary antibodies diluted in blocking buffer without Triton overnight at 4°C. Sections were then washed with PBS and incubated with fluorophore-conjugated secondary antibodies for 1 h at room temperature, washed and mounted using ProLong Gold Anti-fade reagent with DAPI (ThermoFisher Sci, Cat.# P36935) for nuclear stain.

Primary antibodies used: anti-PDGFRα (R&D systems, Cat.# AF1062), anti-EpCAM (Abcam, Cat.# ab71916), anti-GFP (Abcam, Cat.# ab13970), anti-Ki67 (eBioscience, Cat.# 14-5698-82), anti-tdTomato (Takara, Cat.# 632496), anti-K14 (Biolegend, Cat.# 905301), anti-K8 (LSBio, Cat.# LS-B12422), anti-AR (Abcam, Cat.# ab133273) and anti-Laminin (Abcam, Cat.# ab11575). Secondary antibodies used: anti-Goat, anti-Rabbit, anti-Chicken, anti-Rat antibodies conjugated to AlexaFluor 647, AlexaFluor Cy3 and AlexaFluor 488 (Jackson ImmunoResearch). Immunofluorescence staining was imaged using Olympus FLUOVIEW™ FV3000 and FV4000 confocal laser scanning microscopes (Olympus, PA, USA). Images of immunofluorescence staining were analyzed by Fiji ImageJ. Cell number was determined by using the Cell Counter feature in ImageJ.

### Tissue dissociation, flow cytometry and FACS

Immediately after isolation, the entire male urogenital tract was rinsed in PBS on ice. The prostate was isolated from the urogenital tract and placed in warmed digestion solution containing DMEM: F12 with 750 U ml^−1^ collagenase and 250 U ml^−1^ hyaluronidase for 2-2.5 h at 37 °C. Following digestion, digestion mixture was vortexed and centrifuged after addition of Hank’s balanced salt solution with 2% FBS. Digested tissues were treated with ammonium chloride solution to lyse red blood cells, followed by incubation with 0.25% trypsin for 3 min at 37 °C, trituration for another 2 min and then trituration with 5 mg/ml dispase and 0.1 mg/ml DNase for 2 minutes. The resulting solution was then filtered through a 40 µm mesh to obtain a single-cell suspension.

Following dissociation, cells were stained with PE-Cy7 antibodies conjugated to CD45 (Ebioscience, Cat.# 25-0451), CD31 (Ebioscience, Cat.# 25-0311) and Ter119 (Ebioscience, Cat.# 25-5921) to select immune, endothelial, and erythrocytes respectively for exclusion by flow cytometry as the lineage^+^ (Lin^+^) subset. Anti-CD49f/APC (R&D, Cat.# FAB13501A) and anti-EpCAM/APC-Cy7 (Biolegend, Cat.# 118217) were used to identify the lineage^-^ (Lin^-^) stromal and epithelial subpopulations. Dead cells were excluded with Zombie UV viability dye (Biolegend, Cat.# 423107). Endogenous GFP fluorescence was used to detected the GFP-expressing cell population. Flow cytometry was performed using a Fortessa cell analyzer and sorting using a FACSAria Fusion (BD) with FACSDiva software (BD), and FlowJo software was used for downstream analysis (Tree Star, Inc).

### Colony forming cell (CFC) assay

GFP+ and GFP-prostate stromal subpopulations from P35 *Pdgfrα^H2B-eGFP^*mice were FACS-sorted and 300 cells from each population were plated with mitomycin-treated NIH 3T3 fibroblast feeder cells in plates coated with collagen. Colonies were allowed to grow for 10 days, after which cultures were fixed with 4% paraformaldehyde for 15 min and Wright’s Giemsa staining was performed to visualize colonies. Colonies were imaged using an Olympus SZ61 stereo microscope (Olympus, PA, USA)

### Statistical analyses

For all experiments, n represents a distinct biological replicate and data for biological replicates were pooled from independent experiments. Experiments were repeated at least three times with successful replication. Data were analyzed using GraphPad prism software and reported as mean ± standard error of the mean (s.e.m). Comparison of data between two groups was made using Student’s *t*-test (two-tailed). Data between multiple groups were compared using one-way analysis of variance (ANOVA). Statistical significance is recognized at *p*<0.05.

## Acknowledgements

This work was supported by UT Dallas Startup funds to P.A.J. The authors acknowledge the support of Jacob Henderson from the UT Dallas FACS Core for assistance with cell sorting and Joseph Lombardo from the UT Dallas Imaging Core for assistance with confocal microscopy.

## Author contributions

D.M. conceived the study, did bulk of the wet-lab experiments, and wrote the manuscript. H.J.T assisted with wet-lab experiments. P.A.J. conceived, directed the study, and wrote the manuscript with D.M.

## References

1. Aaron, L., Franco, O.E., and Hayward, S.W. (2016). Review of Prostate Anatomy and Embryology and the Etiology of Benign Prostatic Hyperplasia. Urol Clin North Am 43, 279–288. 10.1016/j.ucl.2016.04.012.

2. Baures, M., Dariane, C., Tika, E., Puig Lombardi, E., Barry Delongchamps, N., Blanpain, C., Guidotti, J.E., and Goffin, V. (2022). Prostate luminal progenitor cells: from mouse to human, from health to disease. Nat Rev Urol 19, 201–218. 10.1038/s41585-021-00561-2.

3. Berry, R., and Rodeheffer, M.S. (2013). Characterization of the adipocyte cellular lineage in vivo. Nat Cell Biol 15, 302–308. 10.1038/ncb2696.

4. Brugger, M.D., Valenta, T., Fazilaty, H., Hausmann, G., and Basler, K. (2020). Distinct populations of crypt-associated fibroblasts act as signaling hubs to control colon homeostasis. PLoS Biol 18, e3001032. 10.1371/journal.pbio.3001032.

5. Crowell, P.D., Fox, J.J., Hashimoto, T., Diaz, J.A., Navarro, H.I., Henry, G.H., Feldmar, B.A., Lowe, M.G., Garcia, A.J., Wu, Y.E., et al. (2019). Expansion of Luminal Progenitor Cells in the Aging Mouse and Human Prostate. Cell Rep 28, 1499–1510 e1496. 10.1016/j.celrep.2019.07.007.

6. Cunha, G.R., and Chung, L.W. (1981). Stromal-epithelial interactions--I. Induction of prostatic phenotype in urothelium of testicular feminized (Tfm/y) mice. J Steroid Biochem 14, 1317–1324. 10.1016/0022-4731(81)90338-1.

7. Cunha, G.R., Chung, L.W., Shannon, J.M., Taguchi, O., and Fujii, H. (1983). Hormone-induced morphogenesis and growth: role of mesenchymal-epithelial interactions. Recent Prog Horm Res 39, 559–598. 10.1016/b978-0-12-571139-5.50018-5.

8. Giafaglione, J.M., Crowell, P.D., Delcourt, A.M.L., Hashimoto, T., Ha, S.M., Atmakuri, A., Nunley, N.M., Dang, R.M.A., Tian, M., Diaz, J.A., et al. (2023). Prostate lineage-specific metabolism governs luminal differentiation and response to antiandrogen treatment. Nat Cell Biol 25, 1821–1832. 10.1038/s41556-023-01274-x.

9. Goldstein, A.S., Huang, J., Guo, C., Garraway, I.P., and Witte, O.N. (2010). Identification of a cell of origin for human prostate cancer. Science 329, 568–571. 10.1126/science.1189992.

10. Graham, M.K., Chikarmane, R., Wang, R., Vaghasia, A., Gupta, A., Zheng, Q., Wodu, B., Pan, X., Castagna, N., Liu, J., et al. (2023). Single-cell atlas of epithelial and stromal cell heterogeneity by lobe and strain in the mouse prostate. Prostate 83, 286–303. 10.1002/pros.24460.

11. Ittmann, M. (2018). Anatomy and Histology of the Human and Murine Prostate. Cold Spring Harb Perspect Med 8. 10.1101/cshperspect.a030346.

12. Joshi, P.A., Waterhouse, P.D., Kasaian, K., Fang, H., Gulyaeva, O., Sul, H.S., Boutros, P.C., and Khokha, R. (2019). PDGFRalpha(+) stromal adipocyte progenitors transition into epithelial cells during lobulo-alveologenesis in the murine mammary gland. Nat Commun 10, 1760. 10.1038/s41467-019-09748-z.

13. Kirkwood, P.M., Gibson, D.A., Shaw, I., Dobie, R., Kelepouri, O., Henderson, N.C., and Saunders, P.T.K. (2022). Single-cell RNA sequencing and lineage tracing confirm mesenchyme to epithelial transformation (MET) contributes to repair of the endometrium at menstruation. Elife 11. 10.7554/eLife.77663.

14. Kronstein-Wiedemann, R., and Tonn, T. (2019). Colony Formation: An Assay of Hematopoietic Progenitor Cells. Methods Mol Biol 2017, 29–40. 10.1007/978-1-4939-9574-5_3.

15. Lasnitzki, I., and Mizuno, T. (1980). Prostatic induction: interaction of epithelium and mesenchyme from normal wild-type mice and androgen-insensitive mice with testicular feminization. J Endocrinol 85, 423–428. 10.1677/joe.0.0850423.

16. Lee, Y.H., Petkova, A.P., Mottillo, E.P., and Granneman, J.G. (2012). In vivo identification of bipotential adipocyte progenitors recruited by beta3-adrenoceptor activation and high-fat feeding. Cell Metab 15, 480–491. 10.1016/j.cmet.2012.03.009.

17. Merrick, D., Sakers, A., Irgebay, Z., Okada, C., Calvert, C., Morley, M.P., Percec, I., and Seale, P. (2019). Identification of a mesenchymal progenitor cell hierarchy in adipose tissue. Science 364. 10.1126/science.aav2501.

18. Ousset, M., Van Keymeulen, A., Bouvencourt, G., Sharma, N., Achouri, Y., Simons, B.D., and Blanpain, C. (2012). Multipotent and unipotent progenitors contribute to prostate postnatal development. Nat Cell Biol 14, 1131–1138. 10.1038/ncb2600.

19. Park, J.W., Lee, J.K., Phillips, J.W., Huang, P., Cheng, D., Huang, J., and Witte, O.N. (2016). Prostate epithelial cell of origin determines cancer differentiation state in an organoid transformation assay. Proc Natl Acad Sci U S A 113, 4482–4487. 10.1073/pnas.1603645113.

20. Peng, Y.C., Levine, C.M., Zahid, S., Wilson, E.L., and Joyner, A.L. (2013). Sonic hedgehog signals to multiple prostate stromal stem cells that replenish distinct stromal subtypes during regeneration. Proc Natl Acad Sci U S A 110, 20611–20616. 10.1073/pnas.1315729110.

21. Pignon, J.C., Grisanzio, C., Geng, Y., Song, J., Shivdasani, R.A., and Signoretti, S. (2013). p63-expressing cells are the stem cells of developing prostate, bladder, and colorectal epithelia. Proc Natl Acad Sci U S A 110, 8105–8110. 10.1073/pnas.1221216110.

22. Stingl, J. (2009). Detection and analysis of mammary gland stem cells. J Pathol 217, 229–241. 10.1002/path.2457.

23. Stoyanova, T., Cooper, A.R., Drake, J.M., Liu, X., Armstrong, A.J., Pienta, K.J., Zhang, H., Kohn, D.B., Huang, J., Witte, O.N., and Goldstein, A.S. (2013). Prostate cancer originating in basal cells progresses to adenocarcinoma propagated by luminal-like cells. Proc Natl Acad Sci U S A 110, 20111–20116. 10.1073/pnas.1320565110.

24. Sugimura, Y., Cunha, G.R., and Donjacour, A.A. (1986). Morphogenesis of ductal networks in the mouse prostate. Biol Reprod 34, 961–971. 10.1095/biolreprod34.5.961.

25. Thomson, A.A., and Marker, P.C. (2006). Branching morphogenesis in the prostate gland and seminal vesicles. Differentiation 74, 382–392. 10.1111/j.1432-0436.2006.00101.x.

26. Tika, E., Ousset, M., Dannau, A., and Blanpain, C. (2019). Spatiotemporal regulation of multipotency during prostate development. Development 146. 10.1242/dev.180224.

27. Timms, B.G., Mohs, T.J., and Didio, L.J. (1994). Ductal budding and branching patterns in the developing prostate. J Urol 151, 1427–1432. 10.1016/s0022-5347(17)35273-4.

28. Watson, P.A., Arora, V.K., and Sawyers, C.L. (2015). Emerging mechanisms of resistance to androgen receptor inhibitors in prostate cancer. Nat Rev Cancer 15, 701–711. 10.1038/nrc4016.

29. Wei, X., Zhang, L., Zhang, Y., Cooper, C., Brewer, C., Tsai, C.F., Wang, Y.T., Glaz, M., Wessells, H.B., Que, J., et al. (2022). Ablating Lgr5-expressing prostatic stromal cells activates the ERK-mediated mechanosensory signaling and disrupts prostate tissue homeostasis. Cell Rep 40, 111313. 10.1016/j.celrep.2022.111313.

30. Xie, Q., Liu, Y., Cai, T., Horton, C., Stefanson, J., and Wang, Z.A. (2017). Dissecting cell-type-specific roles of androgen receptor in prostate homeostasis and regeneration through lineage tracing. Nat Commun 8, 14284. 10.1038/ncomms14284.

31. Xin, L., Ide, H., Kim, Y., Dubey, P., and Witte, O.N. (2003). In vivo regeneration of murine prostate from dissociated cell populations of postnatal epithelia and urogenital sinus mesenchyme. Proc Natl Acad Sci U S A 100 Suppl 1, 11896–11903. 10.1073/pnas.1734139100.

