## Supplemental Figures for "PDGFRα^+^ Mesenchymal Stromal Cells Contribute to Epithelial Lineages during Prostate Development"

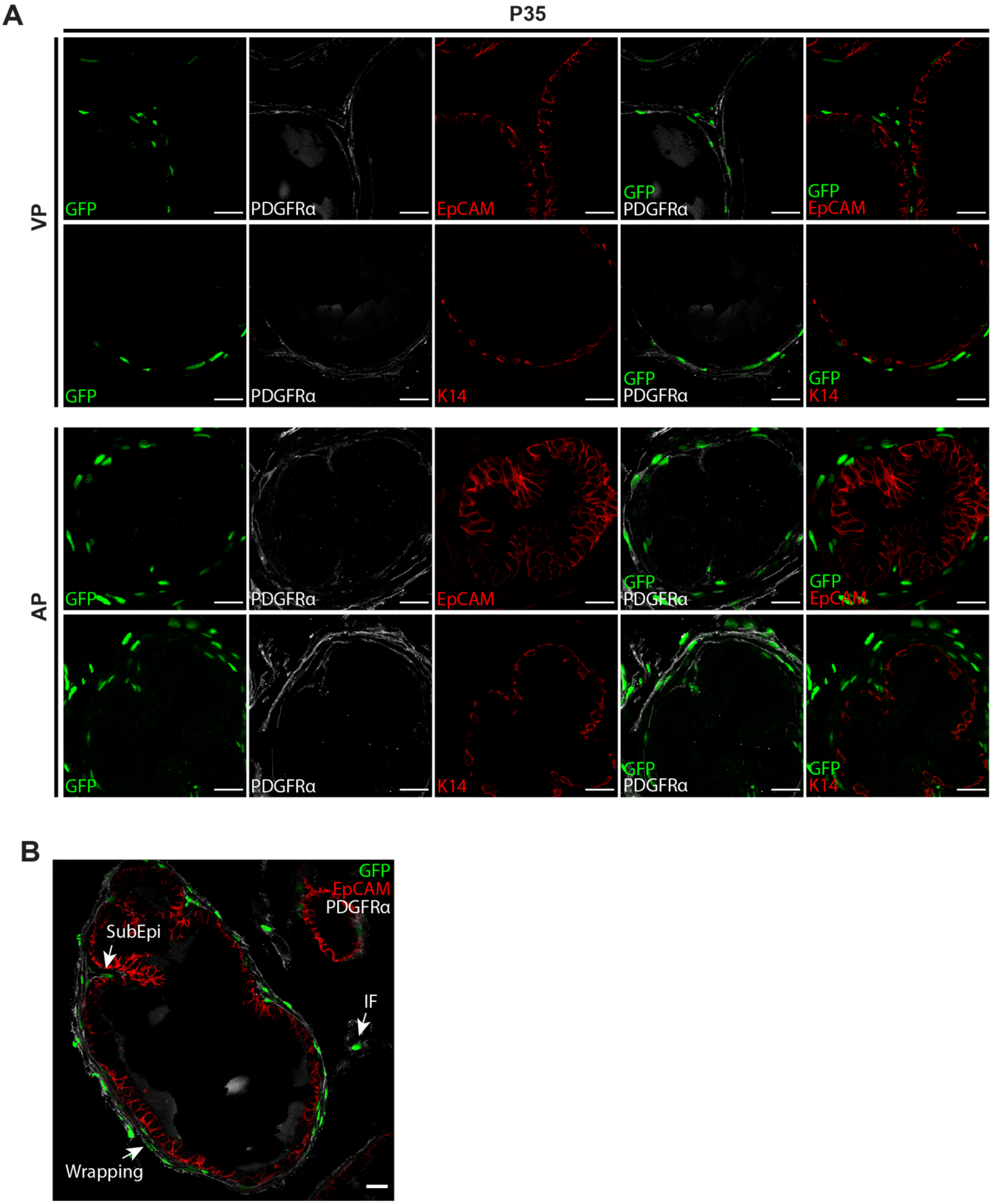
A
P35
VP
AP
GFP
PDGFRα
EpCAM
GFP PDGFRα
GFP EpCAM
K14
GFP K14
B
GFP
EpCAM
PDGFRα
SubEpi
IF
Wrapping

**Figure S1.** ***Pdgfrα*-expressing stromal cells are present in every lobe of the prostate. (A)** Co-immunostaining for GFP, EpCAM, PDGFRα and GFP, K14, PDGFRα in VP and AP lobes of *Pdgfrα*$^{H2B\text{-}eGFP}$ reporter mice at P35 (representative of n = 3 mice each, 3 fields per tissue section), scale bars = 20 µm. **(B)** GFP, EpCAM, PDGFRα in the DLP of reporter mice at P35 showing *Pdgfrα*-expressing GFP$^{+}$ cells in three spatially distinct fibroblast populations: wrapping fibroblasts (Wrapping), subepithelial fibroblasts (SubEpi), and interstitial fibroblasts (IF); scale bars = 20 µm.

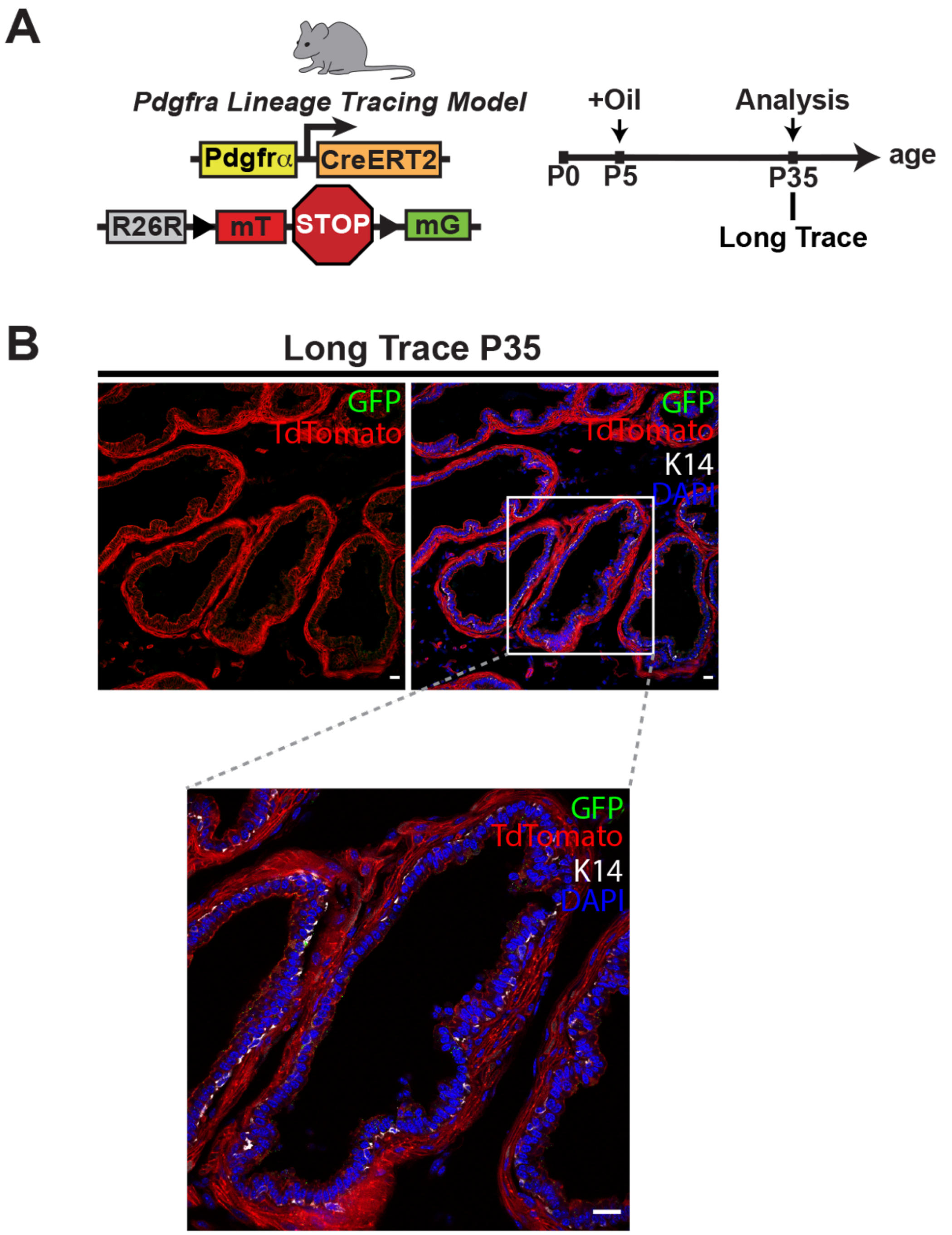


**Figure S2. Lineage tracing control is devoid of GFP-labeled cells. (A)** Schematic of oil (vehicle for TAM) injection and prostate analysis timepoint in *Pdgfrα*$^{CreERT2}$*R26*$^{mTmG}$ lineage tracing mice. **(B)** Immunofluorescence images of GFP, tdTomato and basal epithelial marker K14 in prostate tissue sections from oil-injected controls; scale bars = 20 µm.

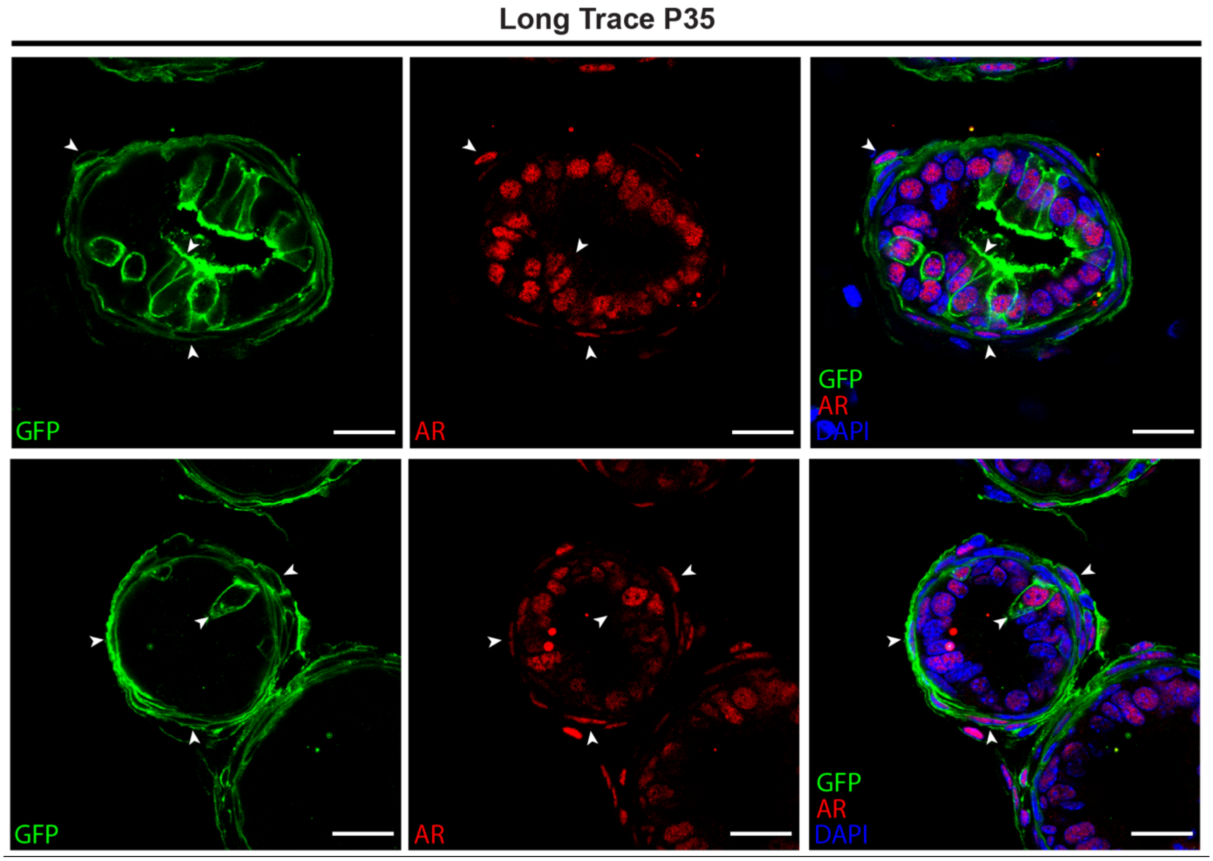


**Figure S3.** ***Pdgfrα*-expressing stromal cells and their epithelial progeny express AR.** Co-immunostaining for GFP with androgen receptor (AR) in the DLP of $\textit{Pdgfr}\alpha^{CreERT2}\textit{R26}^{mTmG}$ lineage tracing mice at P35 (representative of n = 3 mice, 3 fields per tissue section); scale bars = 20 µm. Arrowheads show $GFP^{+}AR^{+}$ stromal and epithelial cells.

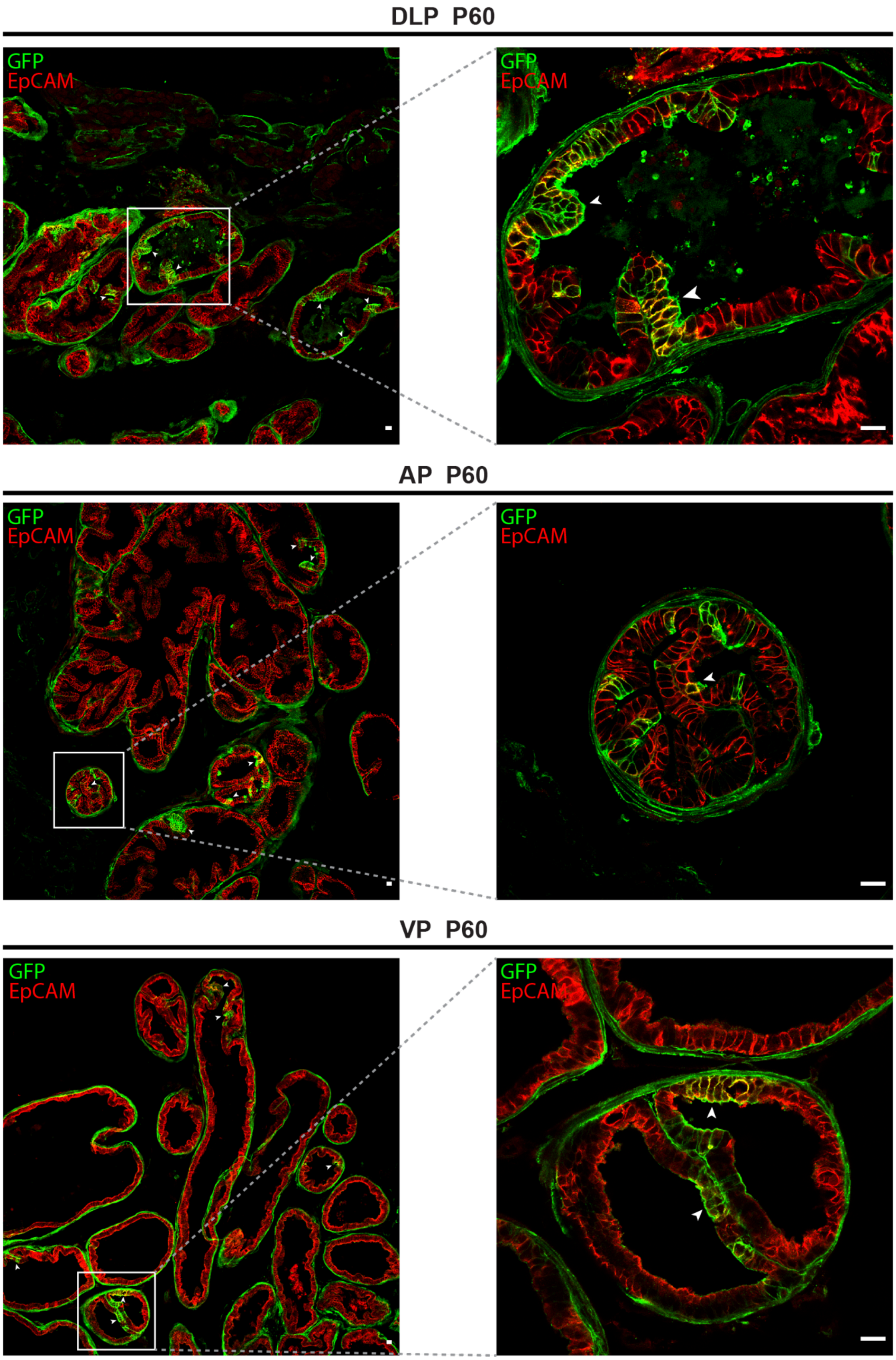
DLP P60
GFP
EpCAM
GFP
EpCAM
AP P60
GFP
EpCAM
GFP
EpCAM
VP P60
GFP
EpCAM
GFP
EpCAM

**Figure S4. Stromal progenitor-derived epithelial cells contribute to every lobe in the mature prostate.**

Co-immunostaining for GFP with epithelial marker EpCAM in the DLP, AP and VP of adult *Pdgfrα*$^{CreERT2}$*R26*$^{mTmG}$ lineage tracing mice at P60 following TAM induction at P5 (representative of n = 3 mice and 3 fields per tissue section); scale bars = 20 µm.
